# Whence Lotka-Volterra? Conservation Laws and Integrable Systems in Ecology

**DOI:** 10.1101/298166

**Authors:** James P. O’Dwyer

## Abstract

Competition in ecology is often modeled in terms of direct, negative effects of one individual on another. An example is logistic growth, modeling the effects of intraspecific competition, while the Lotka-Volterra equations for competition extend this to systems of multiple species, with varying strengths of intra- and inter-specific competition. These equations are a classic and well-used staple of quantitative ecology, providing a framework to understand species interactions, species coexistence, and community assembly. They can be derived from an assumption of random mixing of organisms, and an outcome of each interaction that removes one or more individuals. However, this framing is somewhat unsatisfactory, and ecologists may prefer to think of phenomenological equations for competition as deriving from competition for a set of resources required for growth, which in turn may undergo their own complex dynamics. While it is intuitive that these frameworks are connected, and the connection is well-understood near to equilibria, here we ask the question: when can consumer dynamics alone become an exact description of a full system of consumers and resources? We identify that consumer-resource systems with this property must have some kind of redundancy in the original description, or equivalently there is one or more conservation laws for quantities that do not change with time. Such systems are known in mathematics as integrable systems. We suggest that integrability in consumer-resource dynamics can only arise in cases where each species in an assemblage requires a distinct and unique combination of resources, and even in these cases it is not clear that the resulting dynamics will lead to Lotka-Volterra competition.

I acknowledge the Simons Foundation Grant #376199 and the McDonnell Foundation Grant #220020439.

## 1 Introduction

The phenomenology of competition between species and conditions for their coexistence was formalized over eighty years ago, in a series of works by Lotka, Volterra, Gause, and collaborators [1, 2, 3, 4, 5, 6, during a decade in which progress in quantitative ecology was unprecedented. The result is the familiar set of ordinary differential equations representing competition between species:

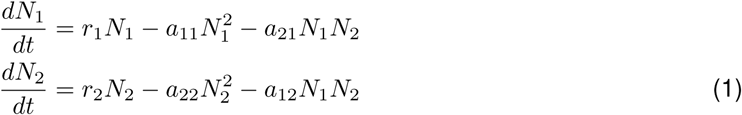

where *r*_1_ and *r*_2_ are per capita growth rates when both species are rare, and the positive parameters *a_ij_* represent intraspecific and interspecific competition. The latter terms are the key signature of the Lotka-Volterra formalism: interactions are represented as quadratic contributions to the rate of change of species densities. These equations can be derived as a deterministic approximation to stochastic processes describing the births and deaths of individuals, where in the case of the quadratic terms the mortality rates are thought of as arising from the ‘collision’ of two individuals in a well-mixed system, and resulting in the removal of one of them [7].

This would be the end of the current paper, except that we as ecologists often prefer to think of these equations as arising from something deeper, and more mechanistic. Even in those seminal papers, the interpretation of the Lotka-Volterra equations was in terms of competition for resources. Gause & Witt [6] explain the different possible dynamics arising from (1) in terms of resources and their availability. For example,

> “When two species belong to different ecological niches in [a] microcosm and, for instance, in addition to common food for each of the species a special kind of food is available that can not be so effectively obtained or consumed by another species, the mutual depression of these species will be less”

In other words, the interpretation was that when the resource overlap for two species is reduced, *a*12 and *a*21 would become smaller. This, and other limits, filled out a qualitative picture for how different values of the parameters *a_ij_* could be motivated in terms of consumer-resource dynamics.

MacArthur and others [8, 9, 10, 11, 12, 13] pioneered efforts to make these early connections between Lotka-Volterra competition and resource consumption quantitative. More recently, contemporary niche theory [14] (based on consumer resource interactions) and modern coexistence theory [12] (phenomenological equations for consumer dynamics alone, like Lotka-Volterra) have made predictions for the properties of equilibria, and whether these equilibria are consistent with coexistence, exclusion, or dependence on initial conditions [15]. But this equilibrium and near-equilibrium analysis leaves a broader question unanswered. Under what circumstances, if any, can phenomenological equations for competition be exactly equivalent to a set of consumer resource dynamics, whether or not the system is close to or far from equilibria?

In this manuscript we address this question. We begin by considering the approach of timescale separation for the consumption of a single resource by a single consumer. Applying this procedure results in a description for consumer dynamics alone, which takes the form of logistic growth. The example illustrates the utility of timescale separation in some regimes, but also demonstrates that it is not guaranteed that the resulting logistic model will be accurate. Moreover, for timescale separation to result in logistic growth for the consumer, it is necessary to assume that the resource dynamics themselves already resemble logistic growth, which is unlikely to hold for the lowest trophic levels, where resources are abiotic. We contrast this with a model of consumers competing for an abiotic resource, and identify a conserved quantity which allows us to eliminate the resources from the model, without assuming a separation of timescales. This resembles previous results in various related fields [16, 17]mathematically as integrabl,mathematically as integrablee and results in exact logistic dynamics for the consumers alone.

We subsequently consider timescale separation for multiple consumers and resources, and illustrate this in a case where the result is Lotka-Volterra competition, Eq.(1). But the same caveats are required as in the case of a single consumer: the dynamics of the resources must take a prescribed form for Lotka-Volterra to arise, and even then there is no guarantee that the Lotka-Volterra description will lead to accurate dynamics away from equilibrium. We therefore extend our previous analysis of a a single abiotic resource to the case of multiple resources. We identify that we need as many conserved quantities as there are distinct resources, in order to ensure an exact description in terms of consumer dynamics alone. Such systems are known mathematically as integrable systems. We present evidence that competition for substitutable resources can not result in an integrable system, while we present a particular model of competition for essential resources, based on multiplicative models of colimitation [18], which does. Intriguingly, the Lotka-Volterra form does not arise in any of the cases we analyze.

## 2 Intraspecific Competition for a Single Resource

### 2.1 Biotic Resource and Separation of Timescales

MacArthur addressed the question of how the full dynamics of a Lotka-Volterra system (and not just its equilibria) might arise from consumer resource dynamics by assuming a separation of timescales [9] between fast resource dynamics and slower consumers. That argument was originally made in the context of (potentially) multiple consumers and multiple, substitutable resources. Here we give an example of how this separation of timescales is used in the case of a single consumer (density *N*) and resource (density *R*). This topic has been extensively revisited [19, 13, 20], for many distinct forms of the underlying dynamics of consumers and resources. In all cases, it is assumed that either there is a separation of timescales between consumer and resource dynamics, or that both consumer and resources are close to their equilibrium densities, or both.

We illustrate this approach using a typical scenario, where biotic resources grow and self-regulate even in the absence of consumers, and consumers only grow through eating resources [21]. We also assume (although this is not essential) that consumers and units of resources meet randomly according to their densities. The result is:

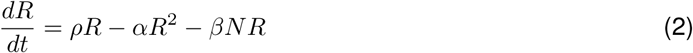

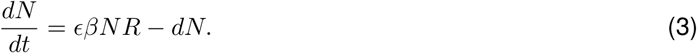

Here, *ρ* and *α* define logistic growth dynamics for the resource in the absence of consumers, while *βNR* is the rate of consumption (with an efficiency *E*). Finally, *d* is an intrinsic mortality rate for consumers. In the case that consumption can be saturated e.g. in the presence of large volumes of resources (and numerous other more realistic assumptions), the terms proportional to *NR* may be replaced by a more general function, *f* (*N, R*).

In its simplest form, the separation of timescales consists of setting 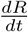 always equal to zero, and then substituting the resulting algebraic constraint into (3). The result sets 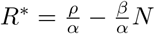 so that:

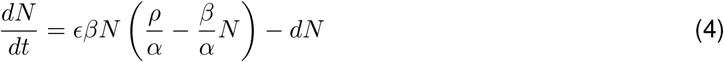

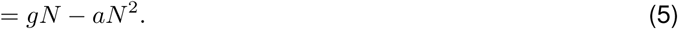

where the effective growth rate 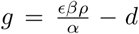, and the effective strength of intraspecific competition is 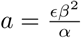.

There are two potentially problematic issues with this approach. The first is: what exactly does a separation of timescales mean here, and how generally can it be valid to replace the dynamics of Eq. (2) with its equilibrium solution? The answer to this question comes from the theory of dynamical systems. The stability properties of an equilibrium can be used to divide the phase space of a dynamical system into qualitatively distinct regions, where these regions are defined with reference to an equilibrium. To put that in ecological terms, the phase space in our example above is the set of all positive consumer and resource densities. For sufficiently small mortality rate, *d*, there is then a single interior equilibrium at

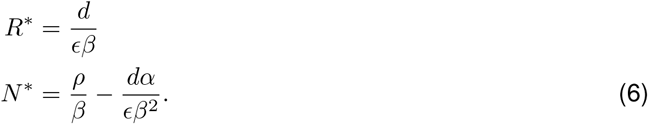

Near to the equilibrium it becomes possible to linearize the system dynamics, and we can divide the phase space up into qualitative regions: the unstable manifold includes all trajectories which diverge from the equilibrium exponentially quickly, the stable manifold all those that converge on the equilibrium exponentially quickly, and the center manifold contains all slower trajectories [22, 23]. This can sometimes greatly simplify dynamics in just the way we wish, as when both stable and center manifolds exist, trajectories will tend to quickly collapse onto the slower dynamics of the center manifold.

Let’s consider this possibility in the context of the equilibrium solution for Eqs. (2) and (3), given by Eq. (6). The Jacobian of the linearized system around its equilbrium is

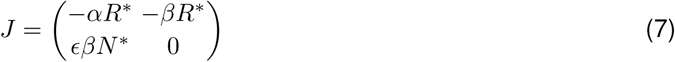

which has eigenvalues

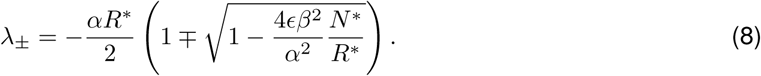

For positive *N ^∗^* and *R^∗^* this equilibrium is therefore locally stable. In fact, there is no center manifold for positive *N ^∗^*, because both eigenvalues are non-zero and hence there is no truly ‘slow’ submanifold onto which trajectories quickly relax. Certainly, there is the potential for faster and slower approaches to the equilibrium, particularly in the limit that 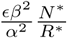 is small. But even in this limit there is no natural division of the slow and fast directions into consumers and resources when both densities are initially far from their equilibrium values. We demonstrate this issue in Figure 1, showing that the logistic approximation to these consumer-resource dynamics can deviate from the approximation of setting the right hand side of Eq. (2) equal to zero, even when there is a reasonable large separation of timescales near to the equilibrium. This contrasts with the dynamics we would have seen, if there were a nontrivial center manifold.

**Fig. 1.**
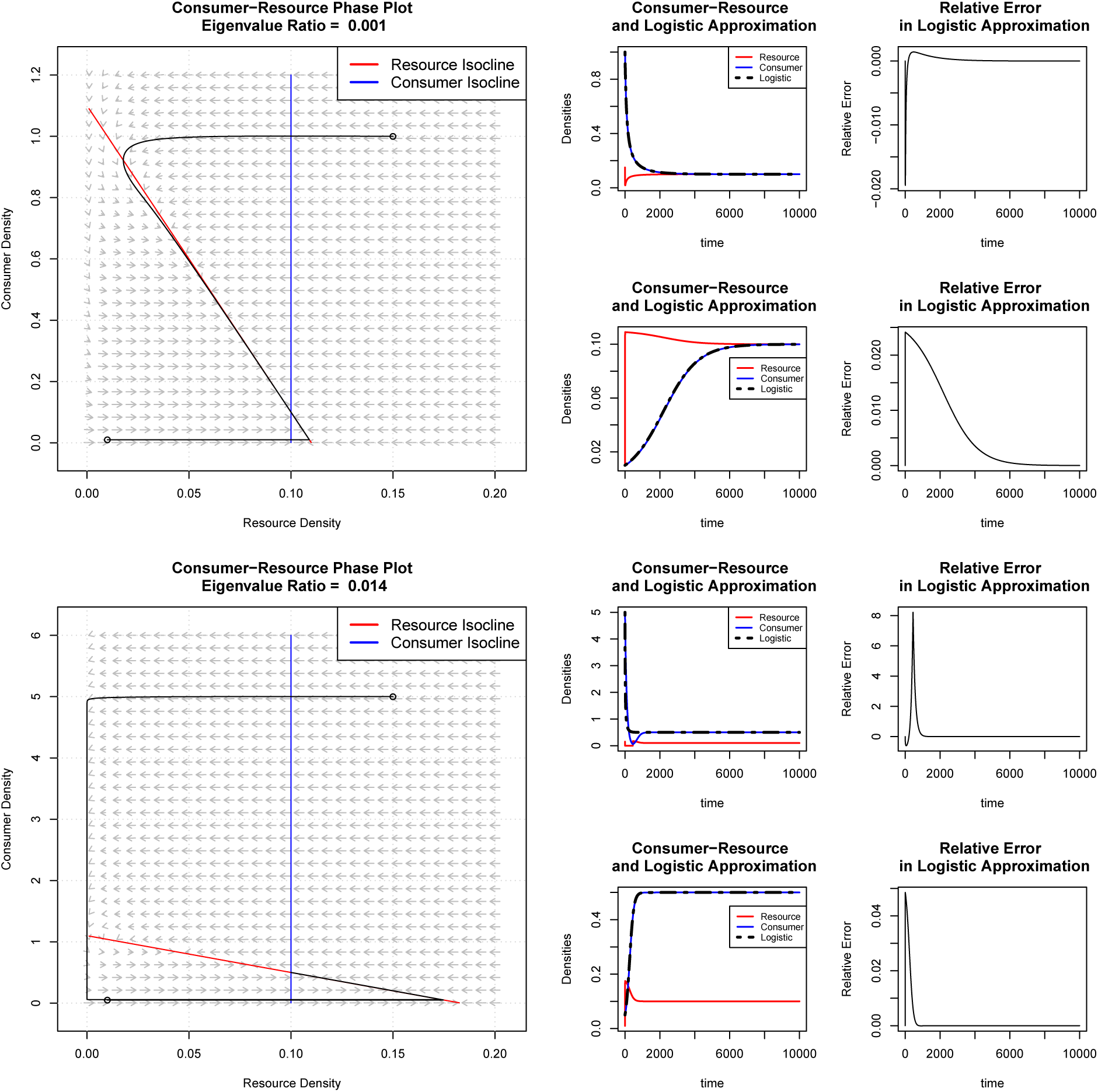
Logistic growth can be a poor approximation to the consumer-resource dynamics of Eqs.(2) and (3), even when there is a separation of timescales near the equilibrium densities for both consumer and resource. We consider two sets of parameter values here in the upper and lower sets of panels—they differ primarily in the ratio of eigenvalues for the linearized system near the equilibrium (where the red and blue nullclines cross), but both have a clear separation of these timescales. For each set of parameter values, two initial conditions are emphasized (black trajectories on the left hand panels). In the first initial condition, both consumer and resource are initially larger than their equilibrium densities, and in the second both are smaller. The corresponding trajectories are plotted in phase space in the left-hand figures, through time in the middle set of figures (for each set of initial conditions), and in the right-hand figures the relative error of using logistic dynamics is plotted. The latter is defined by taking the logistic approximation for consumer density, subtracting the actual consumer density, and dividing the result by consumer density. For the upper set of panels, for a wide range of initial conditions this approximation of setting Eq. (2) to zero is working reasonably well: the dynamics quickly approaches the red nullcline and then proceeds along it. However, it is clear that the initial condition with large values of consumer and resource densities does pass through the red nullcline before returning to it. This feature is exacerbated in the lower set of panels, where the trajectory from this initial condition passes through the red nullcline, deviates substantially from it, before returning to it later on. Even for this (still large) separation of timescales near the equilibrium, we can see substantial deviations from the logistic approximation.

### 2.2 Abiotic Resource and Separation of Timescales

One might think that if most of the dynamics is occurring reasonably close to the equilibrium, the approximation of logistic growth isn’t going to be so bad; indeed, the cases in Figure 1 that go badly wrong are those where we start far from equilibrium. On the other hand, there is a second (and more conceptual) issue with imposing an assumption of separation of timescales, which is also typical in more general cases of timescale separation: we assumed that the resource *R*(*t*) is biotic. In the absence of consumers, Eq. (2) takes the form of logistic growth, and it is this logistic growth that propagates through (via the quadratic consumption terms) to logistic growth for the consumer when we impose a separation of timescales. In other words, there is a kind of fine-tuning here: the form of the equation we get out for the consumers arises from the form of the equation we put in for resources in the first place [20]. For an abiotic variable, intrinsic growth and competition are less reasonable assumptions, and without these we no longer obtain logistic growth for the consumer. For example, suppose that we impose a separation of timescales on the following system:

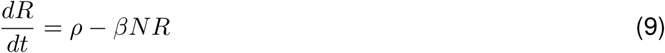

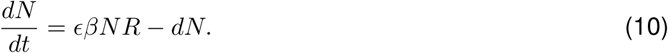

The resulting ODE for consumers is a linear ODE, mirroring the dynamics of the resource [20]. In short, to obtain something as familiar as logistic growth from a separation of timescales, we must assume logistic growth ‘all the way down’ (to the lowest trophic levels). This seems particularly unlikely for the abiotic resources consumed and competed for by microbial communities [24]. In summary, if the lowest trophic levels in an ecological community compete for abiotic resources with an influx from outside the system, then even assuming a separation of timescales does not lead to logistic growth for higher trophic levels.

### 2.3 Abiotic Resource with a Conserved Quantity

Next, we describe a different model of consumer resource dynamics that can be re-expressed as phenomenological dynamics for the species alone. Consider the following system, where a consumer is able to occupy an available abiotic resource, like space:

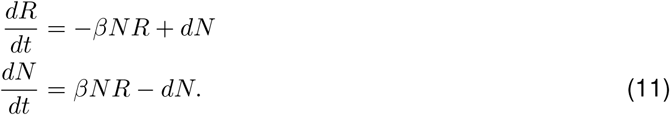

In these equations, consumption of the resource (e.g. occupation of space) is mediated by the same *βNR* term as above, and there is an intrinsic mortality of the consumer, *dN*, which frees up a previously occupied unit of the resource. I am excluding the ‘efficiency’ factor of *E* here by choosing appriopriate units for *R*, so that one unit of the resource (e.g. a region of appropriate spatial area) is equivalent to one consumer. In this system there are two equilibria—either *R* = *d/β* or *N* = 0. There is no separation of timescales, but there is a conserved quantity: *K* = *R* + *N* :

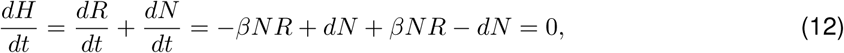

which we could think of as the total available space in this system in the absence of consumers. Then:

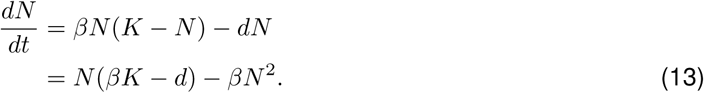

So in this simple example, we have an exact equation for consumers, *N*, which happens to take the form of logistic growth, with a conserved quantity that is interpretable as a carrying capacity. This mirrors a similar elimination of susceptible or infected individuals in the SIS model drawn from epidemiology [17], and Levins’ metapopulation models [16]. It follows even though the resource is interpretable as abiotic (rather than itself undergoing logistic dynamics), and there is no need to make an approximation assuming a separation of timescales. Even more importantly, assuming a separation of timescales by setting 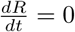 would be a terrible approximation for these consumer-resource dynamics. So at the very least, this approach provides a distinct, parallel derivation of logistic consumer dynamics to that of the previous subsection [17]. In Appendix 1, we go on to relate this to a Hamiltonian formulation, which may become useful in identifying integrable systems with larger numbers of species. We also demonstrate in this Appendix that there are more general systems with a conserved quantity, but where it may still not always be straightforward to use this to integrate out resource dynamics.

## 3 Interspecific Competition among Multiple Consumers

### 3.1 Multiple Biotic Resources and Separation of Timescales

The natural generalization of Eqs. (2) and (3) to the case of two consumers and two substitutable biotic resources is:

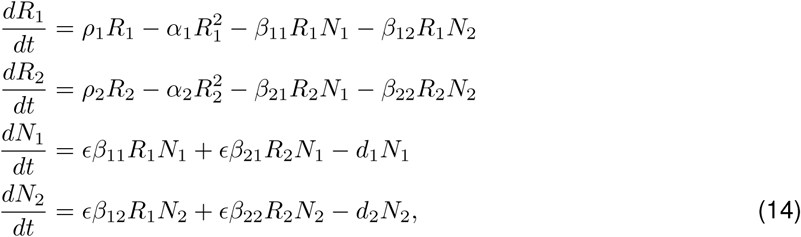

with an obvious generalization to an arbitrary number of each. By applying an assumption of separation of timescales for fast resources, we can rewrite 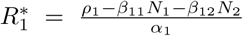, and similarly 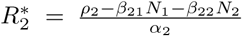. This results in two consumer equations of Lotka-Volterra form:

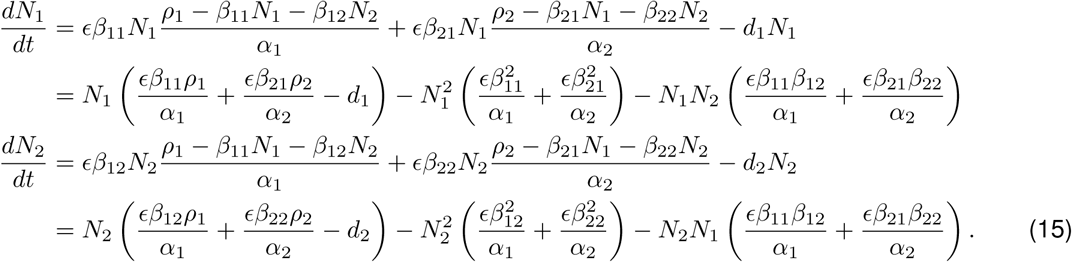

So it is perfectly possible (and already well-known) for us to generalize the separation of time-scales we saw earlier. On the other hand, we are still faced with precisely the same two issues that arise in the case of a single consumer and single resource. First, there is no true slow manifold, and so at best this can be an approximation that will break down when initial conditions are far from equilibrium. Second, there is again a kind of fine tuning of the resource dynamics: the Lotka-Volterra form of the equations we get out critically depends on logistic growth for the biotic resources that we put in.

### 3.2 Substitutable Abiotic Resources

We’ve seen above that consumer-resource dynamics *can* lead to logistic growth for the consumer, without the need to assume either a separation of timescales or logistic dynamics for the resource. This situation arose because there was a conserved quantity and hence a redundancy in the formulation of Eq. (11), in the sense that we can exactly reduce the original description to just a single dynamical variable. A system of *n* resources and *n* consumers would need *n* ‘independent’ conserved quantities in order to (in principle) eliminate all of the explicit resource dynamics exactly, and such cases fall into the realm of integrable dynamical systems.

A system is known as integrable if its phase space (the space of all possible *N_i_* and *R_i_* values) can be foliated by regular invariant submanifolds, where invariance here means that all trajectories initiated within one of these submanifolds remain in that submanifold as time progresses [25]. Put another way, the full phase space can be entirely decomposed into subsets such that if you start within that subset, you are guaranteed to remain within it at all subsequent points in time. The lower the dimension of these invariant submanifolds, the greater the number of equations needed to define their embedding. Each equation takes the general form *f* (*{N_i_}, {R_i_}*) = 0, and hence can be interpreted as a conservation law. This makes clear the meaning of ‘independence’: *m* conserved quantites provide independent constraints if they lead to a well-defined foliation of codimension-*_m_*. For our practical purposes, we will define a consumer-resource system as integrable if we can identify *n* conserved quantities and use them to successfully eliminate *n* resource degrees of freedom from the consumer-resource dynamics.

We now address whether this can be achieved in a system with 2 consumers and 2 substitutable resources. Suppose we have consumers with densities *N*1 and *N*2, and resources with densities *R*1 and *R*2. Is it possible to eliminate both resources, and derive the Lotka-Volterra competition equations for the consumers alone, without imposing a separation of timescales? Perhaps the simplest generalization of the consumer-resource dynamics of Eq. (11) is to allow both consumers to use both of two resources, but at different rates:

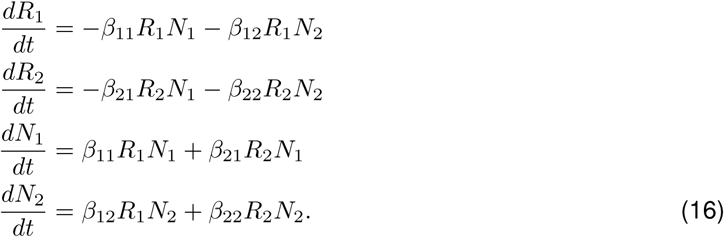

We removed the possibility for mortality of the consumers (and the consequent recycling of consumer biomass into resources), as this does not signficantly change the analysis. These dynamics can be interpreted as a special example of competition for substitutable resources, given that both consumers can survive on either of the two resources (if available). Like the single consumer-single resource system, this set of equations also has a conserved quantity:

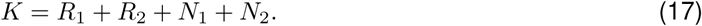

However, this is where the similarity ends. To eliminate both of the resource variables, we would need two conserved quantities. There is no second conserved quantity here, and so it is only possible to eliminate one of the four dynamical variables, not two. Similarly, the generalization of this system of equations to *n* resources and *n* consumers will generically have only one conserved quantity (the exception being special cases where some of the coefficients vanish). In a sense, the reason is obvious. We lose information when a resource is consumed—in contrast to the case of one consumer and one resource, we can no longer directly relate this event to a specific change in the density of a specific consumer. This suggests that while this system of 2 (or more generally *n*) substitutable resources does have a conserved quantity, the dynamics of consumers and substitutable resources can not integrable.

### 3.3 Essential Abiotic Resources and Multiplicative Colimitation

Colimitation of consumers by multiple resources has been extensively investigated in ecology, and is overwhelmingly likely (at some level of description) to be the case in any natural system (see e.g. [26] for examples). While detailed analyses of metabolic networks are (for example in the case of microbial communities) reaching the community level [27, 28, 29], for the most part consumer-resource theory in ecology has focused on tractable phenomenological descriptions of colimitation. The most widely used is known as Liebig’s law [30, 31, 32, 33], where each species is supposed to have a single limiting resource. The resource that is limiting then depends on the environmental context, defined by the concentration or availability of each of a set of essential resources. This is often motivated as a limiting case of a sequence of reactions, where each resource is needed at a given step, but only one step in that sequence is rate limiting. This single step, along with the concentration of the corresponding resource, is (approximately) what determines the reproductive rate of the organism.

Here we consider an alternative model of essential resources, known as multiplicative colimitation [26, 18], where every one of a set of resources determines the growth rate of the organism, in every environmental context. This can be thought of as a situation where an organism must gather all essential resources at the same time, or gain nothing, and leads to a model similar to chemical reaction equations. Both approaches are plausible (if highly coarse-grained) descriptions of the conversion of resources into biomass, and in Appendix B, we show that both Liebig and multiplicative colimitation can arise as approximate limits of the same underlying process. On the other hand, Liebig’s law has often been seen as a better description of colimitation in particular contexts [34] leading Droop [35, 18] to comment that

> “…this argument holds some comfort for ecologists, for it suggests that they may be spared the necessity of becoming biochemists in addition to being mathematicians”

which may resonate with readers of this journal. In any case, there may not be any one definitively correct model of colimitation, and perhaps the detailed biochemistry really does matter in general (flux balance analyses would suggest yes). But for our purposes, multiplicative colimitation will provide a useful example of a plausible integrable system.

Let’s consider an example. Consumer 1 can grow by using 1 unit of resource 1, and *σ* units of resource 2. Consumer 2 can grow by using 1 unit of resource 2, and *τ* units of resource 1. The resulting dynamics are no longer possible to interpret as e.g. occupation of a spatial landscape, as in Eq. (11). In fact, they are identical to chemical reaction equations with appropriate stoichometry, and take the form:

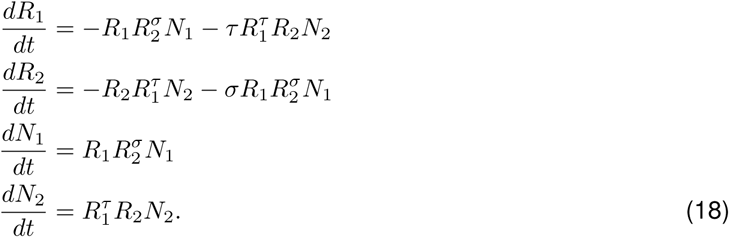

This system now has two conserved quantities:

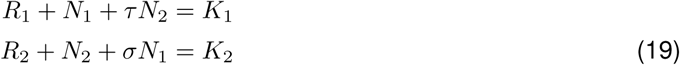

and hence we can exactly write the dynamics of the two species by eliminating the dependencies on *R*_1_ and *R*_2_:

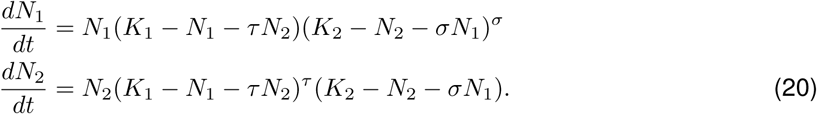

This system of equations is *not* Lotka-Volterra, because it now has higher order terms in consumer densities, that go beyond the quadratic terms in the standard Eq.(1). But such terms could be interpreted as higher-order non-linear interactions between species [40, 36, 37], and so this analysis demonstrates that competition for essential resources provides an example of an integrable set of equations which can be reduced to a reasonable set of dynamics for the consumers alone. What changed when we made resources essential? Because there is a specific (and distinct) combination of the two resources necessary for each consumer to grow, we are able to deduce what must be the changes in *R*_1_ and *R*_2_ if we observe changes in *N*_1_ and *N*_2_ (and vice versa). This was not the case for substitutable resources—by definition, in that case we are unable to deduce from the changes in *N*_1_ and *N*_2_ what specific combination of resource density changes must have occurred. We have not investigated here the parallel case of Liebig’s law colimitation, but it would be interesting to seek more general cases than our multiplicative colimitation example.

This leads us to two speculative conclusions. First, that non-substitutable resources are an essential precondition for integrability, and hence for it to even be possible to write down dynamics for consumers alone. Second, despite the title of our paper, we have not found it possible to obtain precisely Lotka-Volterra equations in *any* model. Perhaps this suggests a challenge for ecologists and mathematicians, or perhaps it suggests that in cases of competition for abiotic resources [24], Lotka-Volterra competition will not be an exact description of consumer dynamics [38, 39].

## 4 Discussion

For almost a century, theoretical ecologists have used density-dependent equations to describe species’ competition for resources. This is exemplified by the Lotka-Volterra competition equations, and is notable for avoiding any specification of the resources for which species compete. This approach, now generalized as modern coexistence theory, is in part pragmatic—sometimes we may observe only species abundances and not resource densities, or we may not know specifically which resources to measure. In parallel, and applied in cases where we do understand something more about resource use and resource dynamics, consumer-resource dynamics (sometimes termed contemporary niche theory [14]) has investigated the dynamics of multiple consumers and resources in tandem. Links between the two frameworks have been surprisingly rare, as reviewed recently [15]. And while it has been possible to rigorously relate the parametrizations of these two bodies of theory near equilibria [15], understanding under what circumstances phenomenological equations for consumer dynamics can exactly recapitulate the dynamics of consumers and resource together is an open question.

We addressed this issue, identifying two problematic issues with the classic approach of assuming a separation of timescales to eliminate resource dynamics from consumer-resource equations. First, this approach can only ever provide an approximation to the consumer dynamics resulting from the full consumer-resource equations. A second issue is that to obtain the specific kinds of consumer dynamics we expect (e.g. logistic growth) it is necessary to assume that the resource dynamics themselves resemble the dynamics of a biotic variable. In cases where resources are abiotic, this is a poor assumption. These issues have already been extensively explored, but to our knowledge not resolved [19, 13, 20].

In this paper we state a simple alternative path: for consumer-resource dynamics to be exactly equivalent to an effective set of dynamics for the consumer alone, there must be some redundancy in the full set of equations. Put simply, if we can identify *n* conserved quantities in a system of *n* consumers and *n* resources, and we can also use these to eliminate the resources from equations for consumer dynamics, then we can reduce the system to a model of consumer dynamics alone. Such dynamical systems are known as integrable, and we identified cases where integrability holds and could be used to integrate out resource dynamics. Specifically, this was possible for a single consumer and single abiotic resource, from which we rederived logistic growth. In combination with the time-scale separation at higher trophic levels, this could provide a clearer justification for logistic growth across the board—but perhaps for different reasons at different trophic levels.

We went on to consider two specific cases of multiple consumers and resources, the kind of situation where one might expect Lotka-Volterra competition to be a reasonable approximation. One could again assume a separation of time-scales in this context, but with the same caveats that there may be both innaccuracy and a certain degree of fine tuning necessary for deriving Lotka-Volterra competition in this way. And so instead we considered consumer-resource dynamics for which there are conserved quantities. From our two cases, we found that only where each consumer had a specific resource requirement, so that resources were essential and not substitutable, was it possible to identify an integrable system with sufficiently-many conserved quantities to exactly integrate out resources. The case of multiple consumers and multiple essential resources we considered is a kind of stoichiometric extreme, where consumers are essentially defined by the specific and unique combination of resources they can convert into new consumers. Its validity might require a fairly radical redefinition of consumers in most contexts.

Does treating these models stochastically, which may be considered a more realistic model of consumption of discrete resources, make the separation of timescales more reasonable? It is possible to make the assumption that the rates determining transitions in resource density are much larger than those leading to transitions in consumer density. But we likely always have some violation of this assumption for certain initial conditions (e.g. initial probability distributions with support for large consumer density). Moreover, we may also have the problem in the case of biotic resources that extinction of some or all of the resources (see also [40]) is the only true equilibrium. So in a sense, the program of deriving consumer dynamics using a separation of timescales in the stochastic versions of these models may be even more challenging than in the deterministic models we considered here.

In summary, to reduce the dynamics of multiple consumers and multiple resources to a simpler set of dynamics for consumers alone, we needed sufficiently many conserved quantities relating each resource separately to the set of consumers. Clearly, these integrable systems are not typical among the set of all possible consumer-resource dynamics, and demanding an exact equivalence between consumer dynamics and the underlying consumer-resource dynamics is fairly strong. On the other hand, it could be that systems that are in some sense close to being integrable still provide a good approximation to this exact description, and so in this sense integrable models may provide a useful baseline. At the very least, our approach gives a parallel derivation of the kinds of consumer dynamics we often use in ecology, which is valid even when separation of timescales is a poor assumption. At the same time, it is interesting that we have not been able to find Lotka-Volterra competition as an exact outcome of this approach (and cannot see a path to doing so), whereas we do find cases leading to higher-order, effective interactions between species [36]. Thus, the title of our paper remains an open question.

## Acknowledgments

I thank Mario Muscarella, Nick Laracuente and Alice Doucet Beaupre´ for constructive criticism on an earlier draft of this manuscript, and Rafael D’Andrea for productive conversations on the history of modeling colimitation. I thank two anonymous reviewers for extremely helpful comments and constructive criticism. I acknowledge Peter Abrams for sending comments and references relating to the separation of timescales approximation, and Sally Otto and Troy Day for commenting on the history of reducing the number of degrees of freedom in SIS and related models.

## A The Hamiltonian Formalism for Consumer-Resource Dynamics

### A.1 Competition for a Single Resource

The construction in the main text for logistic growth (Section 2.2) also forms part of a more general picture. Eqs. (11) are a Hamiltonian system [25, 41], which can be seen by recasting in terms of a pair of canonical variables, say *x*(*t*) and *y*(*t*) and a Hamiltonian, *H*(*x, y*). For a Hamiltonian system, the defining equations must take the form:

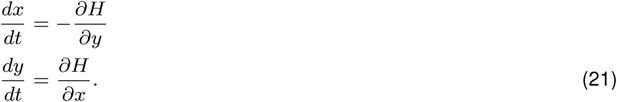

To see this struture explicitly in our case, there is considerable freedom in choosing the variables *x*(*t*) and *y*(*t*). An example is:

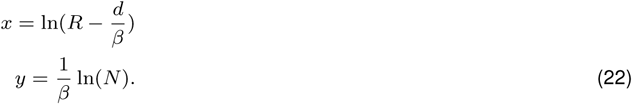

The Hamiltonian is then proportional to the conserved quantity we already saw:

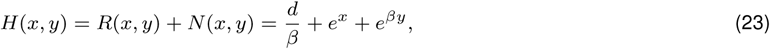

and in terms of these new variables we can rewrite the consumer-resource dynamics explicitly as Hamiltonian dynamics:

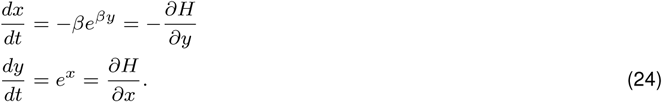

We could also change to another set of canonical variables where the dynamics look even simpler:

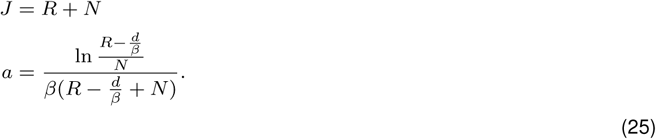

With the Hamiltonian defined as before, *H*(*J, a*) = *J* and hence is independent of the new canonical variable *a*. This leads to very simple equations of motion

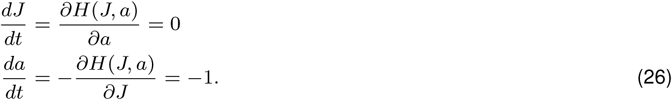

The advantage of these variables is that the solution *a* = *−t* + *const* is straightforward to obtain, and this can of course be translated back into the familiar logistic model solution for *N* (*t*) if necessary.

While the existence of a conserved quantity is on its own sufficient to derive logistic growth for the consumer, making the Hamiltonian formulation slightly superfluous for that specific purpose, we speculate that the structure of Hamiltonian systems may lead to additional insights, beyond this case. This Hamiltonian picture is already known to generalize to a broad range of ecological dynamics [42]. In particular, any two-variable system of the form

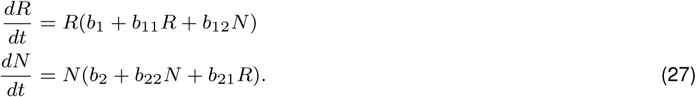

has a conserved quantity and a Hamiltonian formulation provided that *b*_11_*b*_2_(*b*_12_ *− b*_22_) + *b*_22_*b*_1_(*b*_21_ *− b*_11_) = 0 [42]. Our example above (with a redefinition of *R* to *R − d/β*) clearly falls into this category, and another familiar example is the Lotka-Volterra predator-prey equations, for biotic (but unregulated) consumers and resources. This has *b*_11_ = *b*_22_ = 0, and the Hamiltonian

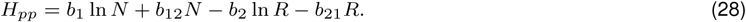

On the other hand, not every Hamiltonian will lead to a single-valued expression for *N* in terms of *R*. For example, if *b*_21_*, b*_12_ *<* 0, and *b*_2_*, b*_1_ *>* 0 in Eq.(28), we cannot rearrange the equation to extract a single-valued solution for *R* in terms of *N* and (the constant) *Hpp*. And so while the existence of a conserved quantity seems essential for finding an exact description of consumer-resource dynamics in terms of consumers alone, it is not necessarily straightforward (or even possible) to use this quantity to derive an ordinary differential equation for *N* alone.

### A.2 Competition for Substitutable Resources

Like the single consumer-single resource system, Equations (16) in the main text can also be recast as a Hamiltonian system, with the Hamiltonian:

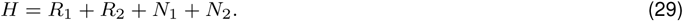

To see the Hamiltonian property explicitly, consider (for example) the change of variables:

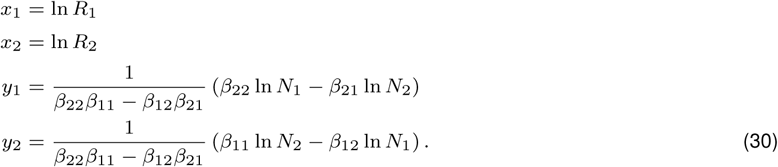

Then with *H*(*x*_1_*, y*_1_*, x*_2_*, y*_2_) = *e^x^*1 + *e^x^*2 + *e^β^*11 *y*1 +*β*21 *y*2 + *e^β^*22 *y*2 +*β*12 *y*1 defined as above

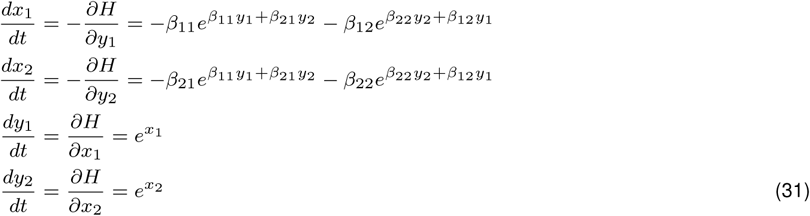

So far, the analysis is very similar to the case of one consumer and one resource. On the other hand, there is no second conserved quantity here, and so it is only possible to eliminate one of the four dynamical variables, not two. Finally, we note that in the case of essential resources considered in Section 3.3, we are unable to find a Hamiltonian formalism.

## B Multiplicative and Liebig Colimitation

In the main text, we motivate a particular form of colimitation by multiple resources, where each resource must be gathered (in a prescribed quantity) at the same time in order for the consumer to add biomass. The resulting equations are essentially the same as chemical reaction equations for chemicals in a vat. We now present a sketch of how both the Liebig form (a single limiting reaction) and multiplicative colimitation can arise from limits of the same sequence of reactions. Consider a consumer, *C* who reaches an intermediate state, *D* after uptake of one unit of resource 1, but where this intermediate state can decay back to state *C*. This is then followed by uptake of resource 2 to make two consumers (i.e. to generate new biomass):

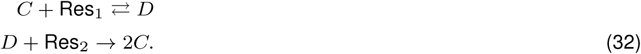

We can represent this system by the ODEs for consumer density *N*, intermediate consumer density *M*, and resource concentrations *R*_1_ and *R*_2_:

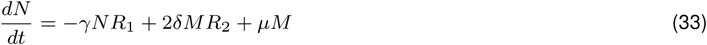

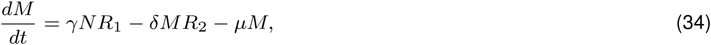

where I am supposing that the resource concentrations are fixed externally or slowly changing, the *γ*, *δ* are various reaction rates and *µ* is a decay rate of the intermediate state back to *C*. Next, suppose that we can assume a separation of timescales (precisely what I’ve argued against in general in the main text, but like those cases, there are limits in which this approximation is reasonable). Then, we can assume 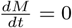 and set

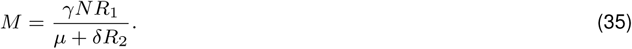

Substituting into Eq. (33), we have that:

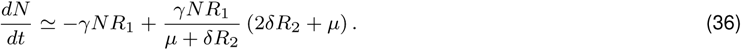

If the decay reaction is rare or absent so that we can ignore *µ*, then this becomes:

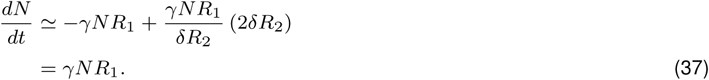

I.ethis limit is consistent with one of the two resources (Res_1_) being rate limiting, and the growth rate of the consumer only depends on this resource. On the other hand, we could also assume that *µ* is very large. In the limit of small 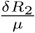,

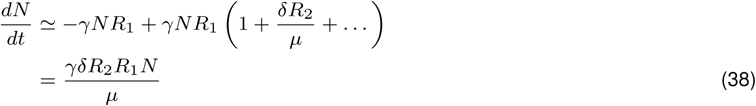

and so we recover the multiplicative form. This is simply saying that if decay via *µ* overwhelms the formation of the intermediate state, *D*, then in effect the consumer still needs to gather both resources simultaneously. Finally, if we assume a separation of timescales but now assume that the first reaction is fast, then we can approximately set 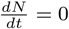 and obtain

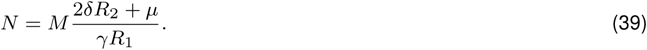

Substituting this into the remaining equation we have

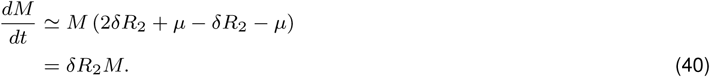

Hence both the intermediate state, *D*, and by Eq. (39) the consumer *C*, grow exponentially with rate determined by *δR*_2_. I.e. the second resource is limiting, rather than the first. Therefore, with the caveats necessary to trust this ‘separation of timescales’ approach, all three cases (both of Liebig’s possible laws of the minimum, and multiplicative colimitation) can arise as limiting cases of this simple sequence of reactions. It’s unclear whether either approximation dominates in colimitation in nature, though there seems to be stronger evidence for Liebig. Perhaps a more systematic approximation method for microbial metabolic networks may pay dividends in understanding which of these cases (if any) apply in a given natural system.

